# Gut microbiota and resistome dynamics in intensive care patients receiving selective digestive tract decontamination

**DOI:** 10.1101/102467

**Authors:** Elena Buelow, Teresita de Jesús Bello González, Susana Fuentes, Wouter A.A. de Steenhuijsen Piters, Leo Lahti, Jumamurat R. Bayjanov, Eline A.M. Majoor, Johanna C. Braat, Maaike S. M. van Mourik, Evelien A.N. Oostdijk, Rob J.L. Willems, Marc. J.M. Bonten, Mark W.J. van Passel, Hauke Smidt, Willem van Schaik

**Affiliations:** Department of Medical Microbiology, University Medical Center Utrecht, Utrecht, The Netherlands; Laboratory of Microbiology, Wageningen University, Wageningen, The Netherlands; Department of Pediatric Immunology and Infectious Diseases, The Wilhelmina Children’s Hospital, University Medical Center Utrecht, Utrecht, The Netherlands; Department of Mathematics and Statistics, University of Turku, Turku, Finland.; Center of Infectious Disease Control, National Institute of Public Health and the Environment, Bilthoven, The Netherlands

**Keywords:** Anti-Bacterial Agents, Antibiotic Prophylaxis, Drug Resistance, Microbial, Intensive Care, Microbiome

## Abstract

**Background:** Critically ill patients hospitalized in an Intensive Care Unit (ICU) are at increased risk of acquiring potentially life-threatening infections with opportunistic pathogens. The gut microbiota of ICU patients forms an important reservoir for these infectious agents. To suppress gut colonization with opportunistic pathogens, a prophylactic antibiotic regimen, termed ‘Selective decontamination of the digestive tract’ (SDD), may be used. SDD has previously been shown to improve clinical outcome in ICU patients, but the impact of ICU hospitalization and SDD on the gut microbiota remains largely unknown. Here, we characterize the composition of the gut microbiota and its antimicrobial resistance genes (‘the resistome’) of ICU patients during SDD.

**Results:** During ICU-stay, 30 fecal samples of ten patients were collected. Additionally, feces were collected from five of these patients after transfer to a medium-care ward and cessation of SDD. As a control group, feces from ten healthy subjects were collected twice, with a one-year interval. Gut microbiota and resistome composition were determined using 16S rRNA phylogenetic profiling and nanolitre-scale quantitative PCRs.

The microbiota of the ICU patients differed from the microbiota of healthy subjects and was characterized by low microbial diversity, decreased levels of *E. coli* and of anaerobic Gram-positive, butyrate-producing bacteria of the *Clostridium* clusters IV and XIVa, and an increased abundance of Bacteroidetes and enterococci. Four resistance genes (*aac(6′)-Ii*, *ermC*, *qacA*, *tetQ*), providing resistance to aminoglycosides, macrolides, disinfectants and tetracyclines respectively, were significantly more abundant among ICU patients than in healthy subjects, while a chloramphenicol resistance gene (*catA*) and a tetracycline resistance gene (*tetW*) were more abundant in healthy subjects.

**Conclusions:** The microbiota and resistome of ICU patients and healthy subjects were noticeably different, but importantly, levels of *E. coli* remained low during ICU hospitalization, presumably due to SDD therapy. Selection for four antibiotic resistance genes was observed, but none of these are of particular concern as they do not contribute to clinically relevant resistance. Our data support the ecological safety of SDD, at least in settings with low levels of circulating antibiotic resistance.

## Background

The human gut microbiota comprises 10^13^ - 10^14^ bacterial cells that belong to hundreds of different species. The gut microbiota plays an important role in numerous metabolic, physiological, nutritional and immunological processes of the human host [1]. In healthy individuals, the gut microbiota mostly consists of bacteria that have a commensal or mutualistic relationship with the human host. Critically patients, however, frequently have an extremely dysbiotic gut microbiota that is characterized by intestinal overgrowth with multi-drug resistant opportunistic pathogens of the phylum Proteobacteria (e.g. *Escherichia coli*) and the genus *Enterococcus*, while the abundance of commensal *Bacteroidetes* and *Firmicutes* is decreased [2–5]. The high levels of aerobic, opportunistic pathogens in the gut during critical illness are likely contributing to the burden of respiratory and bloodstream infections with these organisms in critically ill patients [6]. Selective Digestive tract Decontamination (SDD) aims to reduce the risk of nosocomial infections in ICU patients. SDD aims to eradicate opportunistic pathogens from the patients, while minimally impacting commensal bacteria [7]. In SDD a paste containing the antibiotics colistin and tobramycin, and the antifungal amphotericin B, is applied to the oropharynx of ICU patients. The patients also receive a suspension of colistin, tobramycin, and amphotericin B via a nasogastric tube. These antimicrobials are applied from the day of ICU admission until ICU discharge. In addition, a third-generation cephalosporin (usually either cefotaxime or ceftriaxone) is administered intravenously during the first 4 days of ICU stay. SDD lowers patient mortality during ICU stay in settings with a low prevalence of antibiotic resistance and reduces the costs associated with ICU hospitalization [8,9]. Selection of bacteria that are resistant to the antimicrobials used in SDD remains a major concern [10,11], although this is not supported by the results of clinical trials in which conventional culture techniques were used to screen for antibiotic resistance among nosocomial pathogens [12]. The patient gut is not only a potential source for opportunistic pathogens, but also forms a large reservoir for antibiotic resistance genes, termed the gut resistome [13–17]. The use of antibiotics may favor the selection for antimicrobial resistance genes (ARGs) among members of the gut microbiota, thus increasing the likelihood of horizontal spread of ARGs between commensals and opportunistic pathogens co-residing in the gut [16]. During the administration of SDD, the gut resistome of patients is monitored by the cultivation of resistant bacteria from rectal swabs or feces, as part of routine diagnostics. However, methods that rely on microbial culture capture only a fraction of the gut resistome, since anaerobic commensals, which are the main reservoir of ARGs in the gut microbiota, are difficult to culture [18–20]. Thus, culture-independent methods are needed to comprehensively assess the impact of antibiotic prophylaxis on the microbiota and resistome of ICU patients.

Here, we used the 16S ribosomal RNA (rRNA) gene-targeted Human Intestinal Tract Chip (HITChip) and nanolitre-scale quantitative PCR (qPCR) targeting 81 ARGs, to determine the dynamics of gut microbiota composition and resistome in patients receiving SDD during ICU hospitalization. We contrast these findings in ICU patients with the composition of the microbiota and resistome of healthy subjects.

## Methods

### Study population

All included patients (*n* = 10) were acutely admitted to the ICU of the University Medical Center Utrecht from the community and had not been hospitalized in the previous six months, with the exception of patient 105 who was hospitalized for 5 days prior to transfer to the ICU and start of SDD. None of the patients were treated with antibiotics in six months prior to ICU hospitalization. All patients received SDD from the start of ICU stay until ICU discharge. SDD consists of 1000 mg of cefotaxime intravenously four times daily for four days, an oropharyngeal paste containing polymyxin E, tobramycin and amphotericin B (each in a 2% concentration) and administration of a 10 mL suspension containing 100 mg polymyxin E, 80 mg tobramycin and 500 mg amphotericin B via a nasogastric tube, four to eight times daily throughout ICU stay. All patients received additional antibiotics during ICU stay. Fecal samples of patients were collected by nursing staff upon defecation and stored at 4°C for 30 min to 4 h, after which the samples were transferred to -80°C. Seven patients included here (patient numbers: 105, 108, 120, 163, 164, 165 and 169) were also included in a previous study where the dynamics of two aminoglycoside resistance genes in the gut microbiota of ICU patients was studied [20]. A total of 30 fecal samples were collected during ICU stay. Five additional fecal samples were collected after transfer to a medium care ward and cessation of SDD. Fig. S1 includes detailed information on sampling time points and antibiotic usage of the ICU patients in this study.

Routine surveillance for colonization with aerobic Gram-negative bacteria in ICU patients was performed through culturing of rectal swabs on sheep blood agar and MacConkey agar. All suspected Gram-negative colonies were analyzed by Maldi-TOF for species identification. Antibiotic resistance phenotypes were determined using the Phoenix system (BD, Franklin Lakes, NJ, USA). Fecal samples of healthy subjects were collected as part of the ‘Cohort study of intestinal microbiome among Irritable Bowel Syndrome patients and healthy individuals’ (CO-MIC) study at two time-points with a one-year interval between sampling. None of the individuals in this cohort received antibiotics. All included patients and healthy subjects were adults.

### Gut microbiota profiling by HITChip

The HITChip is a validated phylogenetic array produced by Agilent Technologies (Palo Alto, CA) and developed at Wageningen University, The Netherlands [21,22]. It contains over 4,800 oligonucleotides targeting the V1 and the V6 region of the 16S rRNA gene from 1,132 microbial phylotypes present in the human gut [21]. DNA from fecal samples was isolated as previously described [23]. The full-length 16S rRNA gene was amplified from fecal DNA, and PCR products were further processed and hybridized to the microarrays as described previously [24]. Data analyses were performed using R (www.r-project.org), including the microbiome package (https://github.com/microbiome). Bacterial associations in the different patient groups and healthy subjects were assessed using Principal Component Analysis (PCA) as implemented in CANOCO 5.0 [25]. Statistical testing for the differences in microbiota composition between ICU patients and healthy subjects was performed by the non-parametric Mann-Whitney U test. All P-values were corrected for false discovery rate (FDR) by the Benjamini and Hochberg method [26], and corrected P-values (*q*) below 0.05 were considered significant.

### Quantification of *E. coli* by qPCR

QPCRs for the quantification of *E. coli* were performed with primers that were previously described [27], using serial dilutions of genomic DNA of *E. coli* DH5α to generate a standard curve. The quantification of 16S rRNA was performed with primers described in [28]. The PCR conditions were identical to the qPCR conditions for the detection of *mcr-1* (described above).

### qPCR analysis of antibiotic resistance genes

qPCR analysis was performed using the 96.96 BioMark™ Dynamic Array for Real-Time PCR (Fluidigm Corporation, San Francisco, CA), according to the manufacturer’s instructions, with the exception that the annealing step in the PCR was held at 56°C. Fecal DNA was first subjected to 14 cycles of Specific Target Amplification using a 0.18 μM mixture of all primer sets, excluding the 16S rRNA primer sets, in combination with the Taqman PreAmp Master Mix (Applied Biosystems), followed by a 5-fold dilution prior to loading samples onto the Biomark array for qPCR. Thermal cycling and real-time imaging was performed on the BioMark instrument, and Ct values were extracted using the BioMark Real-Time PCR analysis software.

### Target selection, primer design and primer validation

The primer set used in the qPCR assays covered 81 antimicrobial resistance genes (ARGs) of 14 resistance gene classes (Table S1). Primers were designed for the ARGs that are most commonly detected in the gut microbiota of healthy individuals [14,15] and clinically relevant ARGs, including genes encoding extended spectrum β-lactamases (ESBLs), carbapenemases, and proteins involved in vancomycin resistance. We also included 10 genes encoding transposases, and a gene encoding an integrase as representatives of mobile genetic elements [29]. Seven of these genes were detected by qPCR but no significant differences could be observed between patients and healthy subjects (data not shown) and these are not further discussed in our manuscript. Primer design was performed using Primer3 [30] with its standard settings with a product size of 80 – 120 bp and a primer melting temperature of 60°C. The universal primers for 16S rRNA genes were previously described by Gloor *et al.* [28]. Forward and reverse primers were evaluated *in silico* for cross hybridization using BLAST [31] and were cross-referenced against ResFinder [32] to ensure the correct identity of the targeted genes. All primers that aligned with more than 10 nucleotides at their 3’ end to another primer sequence were discarded and redesigned. Additionally, all primer sets were aligned to all resistance genes that were targeted in this PCR analysis to test for cross hybridisation with genes other than the intended target resistance gene. Primers that aligned with more than 10 nucleotides at their 3’ end sequence with a non-target resistance gene were discarded and redesigned. A reference sample consisting of pooled fecal DNA from different patients was loaded in a series of 4-fold dilutions and was used for the calculation of primer efficiency. All primers whose efficiency was experimentally determined to be between 80% and 120% were used to determine the normalized abundance of the target genes. The detection limit on the Biomark system was set to a CT value of 20, as recommended by the manufacturer. In addition, to assess primer specificity we performed melt curve analysis using the Fluidigm melting curve analysis software (http://fluidigm-melting-curve-analysis.software.informer.com/). All PCRs were performed in triplicate and sample-primer combinations were only included in the analysis when all triplicate reactions resulted in a CT-value below the detection limit.

After completion of the nanolitre-scale qPCR assays, the transferable colistin resistance gene *mcr-1* was described [33]. To detect and quantify *mcr-1*, we developed primers (qPCR-mcr1-F: 5’-TCGGACTCAAAAGGCGTGAT-3’ and qPCR-mcr1-R: 5’-GACATCGCGGCATTCGTTAT-3’) for use in a standard qPCR assay. The *mcr-1* gene was synthesized based on the sequence described in [33] by Integrated DNA Technologies (Leuven, Belgium) and used as a positive control in our assays. The qPCR was performed using Maxima SYBR Green/ROX qPCR Master Mix (Thermo Scientific, Leusden, The Netherlands) and a StepOnePlus instrument (Applied Biosystems, Nieuwekerk a/d IJssel, The Netherlands) with 5 ng DNA in the reaction and the following program: 95°C for 10 min, and subsequently 40 cycles of 95°C for 15 sec, 56°C for 1 min.

### Calculation of normalized and cumulative abundance

Normalized abundance of resistance genes was calculated relative to the abundance of the 16S rRNA gene (CT_ARG_ − CT_16S rRNA_), resulting in a log2-transformed estimate of ARG abundance. Statistical analysis was performed using GraphPad Prism (La Jolla, CA). Visualization of the qPCR data in the form of a heat map was performed using Microsoft Excel. Statistical testing of the differences in the abundance of resistance genes in ICU patients versus healthy subjects was performed with the non-parametric Mann-Whitney U test with a False Discovery Rate (Benjamini Hochberg) < 0.05 to correct for multiple testing.

## Results

### Microbiota dynamics in ICU patients and healthy subjects

Global changes in the gut microbiota of healthy subjects and ICU patients were visualized by Principal Component Analysis (Fig. 1A). The microbiota profiles of healthy subjects clustered together, indicating that they had stable and broadly comparable microbiota profiles, which were clearly distinct from the microbiota profiles of patients during and after ICU stay. These profiles covered a larger area in the PCA plot, indicating that the differences in the microbiota composition of patients were larger than in healthy subjects. Notably, the microbiota of several ICU patients (e.g. #120, #169, #180, #43) was already distinct from the microbiota of the healthy subjects in the first days of ICU hospitalization, indicating that ICU hospitalization has a rapid effect on the microbiota. The diversity of the microbiota, as quantified by Shannon’s diversity index, was significantly lower in ICU patients compared to healthy subjects (5.90 ± 0.20 vs 5.19 ± 0.46, respectively; *P* < 0.001, Student’s t-test). The diversity of the microbiota of ICU patients was highly dynamic (Fig. 1B). Several patients (#108, #163, #164, #165 and #169) experienced a rapid loss of diversity in the first days of ICU stay. In contrast, the diversity of the microbiota was more stable in healthy subjects when comparing samples that were collected one year apart (Fig. 1C). Compared to healthy subjects, the microbiota of patients during ICU hospitalization was characterized by a significantly higher abundance in the taxa Bacteroidetes and Bacilli: *Enterococcus* and a lower abundance of the taxa *Clostridium* cluster IV and XIVa (Fig. 1D).

**Figure 1.**
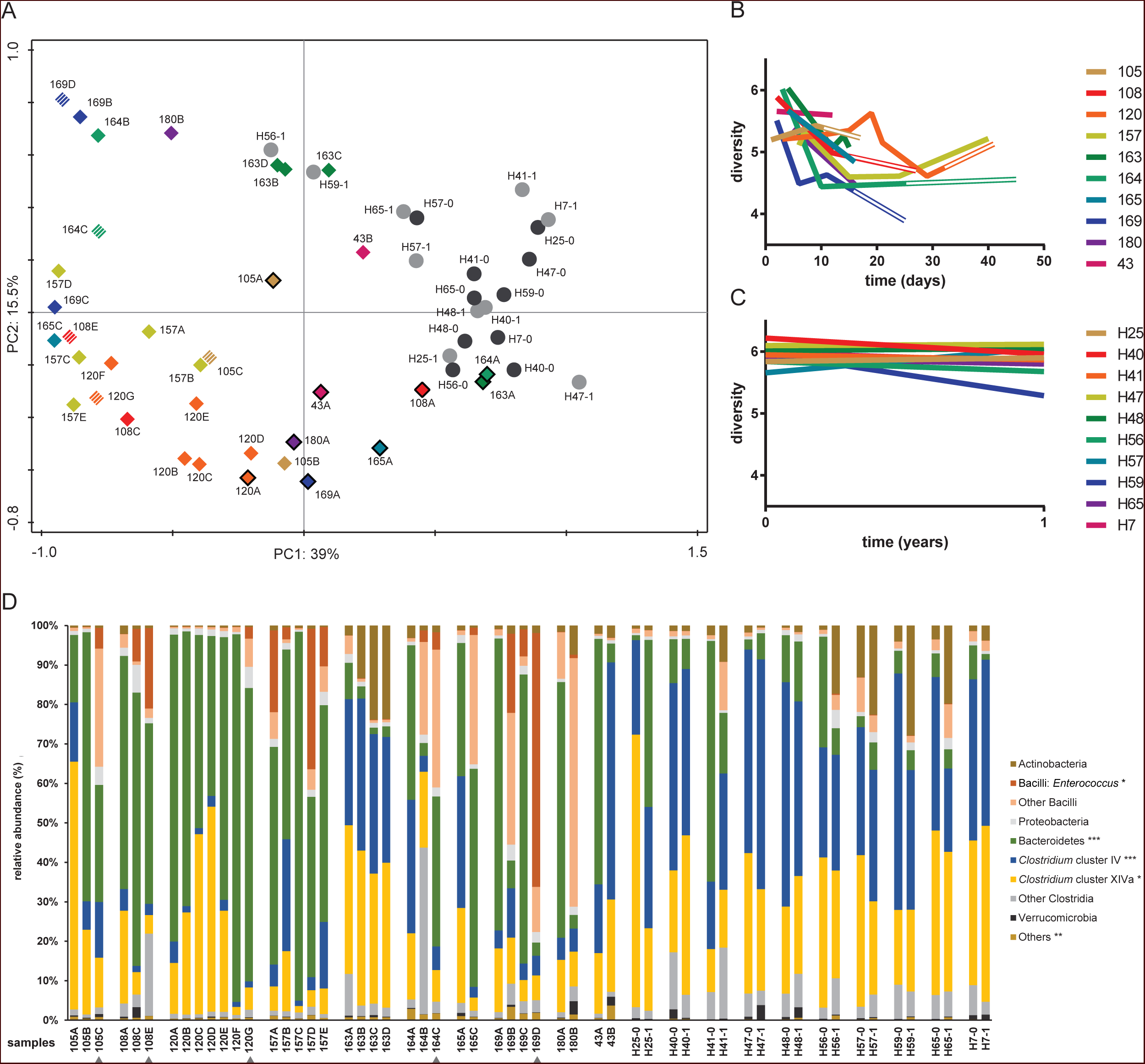
Dynamics of gut microbiota composition and diversity in ICU patients and healthy subjects. (A) Principal Component Analysis (PCA) of gut microbiota composition of ICU patients.Dashed symbols indicate fecal samples collected after ICU discharge and continued hospitalization in a medium-care ward. Fecal samples that were collected in the first 5 days of ICU hospitalization are indicated by a black line around the symbol. Fecal samples of healthy subjects were collected at two time-points with one-year interval, indicated with black and grey circles, respectively. (B) Diversity (Shannon index) of the microbiota of ICU patients. Double lines indicate hospitalization in a medium-care ward. (C) Diversity (Shannon index) of the microbiota of healthy subjects. (D) Gut microbiota composition of patients and healthy subjects. Stacked bar charts represent the abundance of different major taxa in the gut microbiota of ICU patients and healthy subjects. Among Bacilli, the genus *Enterococcus* has been highlighted, as SDD has previously been shown to select for colonization with enterococci [38,39]. Fecal samples that were collected after ICU discharge and during medium-care hospitalization are indicated by grey triangles. Statistically significant differences of the abundance of taxa in the gut microbiota of patients during ICU hospitalization and healthy subjects are indicated in the legend (*: *q* < 0.05; **: *q* < 0.01; ***: *q* < 0.001; Mann-Whitney U test with Benjamini-Hochberg correction for multiple testing).

We performed quantitative PCRs to accurately determine the abundance of *E. coli*, one of the primary targets of SDD, in the gut microbiota of patients and healthy subjects (Fig. 2). The abundance of *E. coli* in samples of ICU patients was lower compared to the healthy subjects (*p* = 0.001; Mann-Whitney U test). Notably, upon cessation of SDD and transfer to a medium-care ward, the abundance of *E. coli* rebounded in one patient (#105) to levels surpassing those found in healthy individuals.

**Figure 2.**
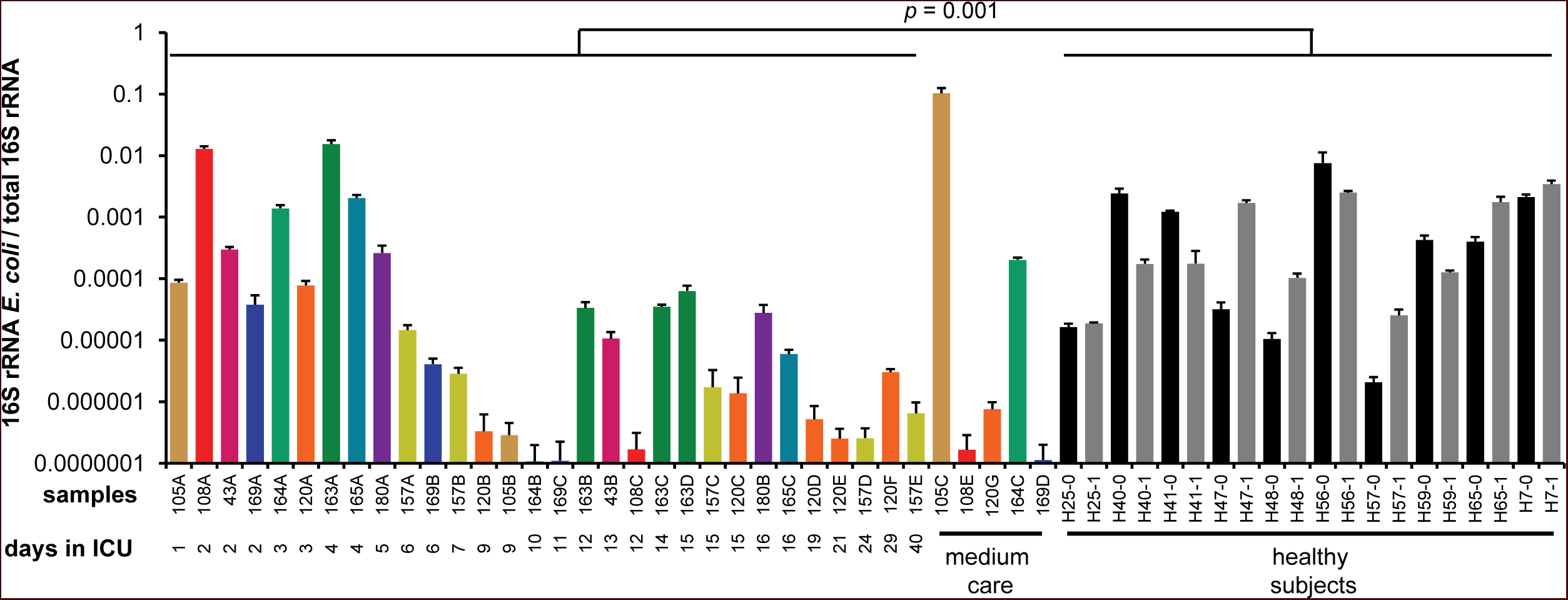
Abundance of *E. coli* in the gut microbiota of ICU patients and healthy subjects. Quantification of *E. coli* 16S rRNA gene copies relative to total 16S rRNA gene copies, performed by qPCR with three technical replicates. Error bars indicate standard deviation. Samples are ordered by time of sampling during ICU stay. The color coding of the samples is unique for each patient and is identical to Fig 1. Statistical testing was performed with the Mann-Whitney U test.

During ICU stay, routine surveillance by conventional microbiological culture was performed on all patients. *E. coli* could be cultured from six out of 73 rectal swabs that were collected during the patients’ ICU-stay. Five *E. coli* positive rectal swabs, of patients #43, #105, #108, #163, and #169, were collected within one day of ICU admission, while the sixth positive swab (of patient #165) was collected after nine days of ICU-stay. In addition, an *E. coli* strain from patient #105 with an ESBL-producing and tobramycin-resistant phenotype was isolated after ICU discharge, while the patient was in a medium-care ward. The *E. coli* strains isolated during ICU stay were susceptible to cephalosporins and aminoglycosides. All *E. coli* strains were susceptible to colistin.

### Resistome dynamics in ICU patients and healthy subjects

A total of 46 unique ARGs conferring resistance to 12 different classes of antimicrobials were detected in the DNA isolated from fecal samples of hospitalized patients and healthy subjects (Fig. S2). The number of detected resistance genes per sample ranged between 6 and 38. Eleven resistance genes were detected in >80% of healthy subjects and critically ill patients. This highly prevalent set of resistance genes included tetracycline resistance genes (*tetO, tetQ*, *tetM*, *tetW*), two aminoglycoside resistance genes (*aph(3′)-III* and an *aadE-*like gene), the bacteroidal β-lactam resistance gene *cblA*, and the macrolide resistance gene *ermB*.

Genes associated with major antibiotic resistance threats, including those identified by the Centers for Disease Control, were relatively rare. Genes encoding for extended-spectrum beta-lactamases (ESBLs) were not detected in healthy subjects. In two ICU patient samples (#105C and #108C), however, the ESBL genes *bla_CTX-M_* and *bla_DHA_*, respectively, could be detected. Sample #105C was collected after ICU discharge and cessation of SDD, while sample #108C was collected after 12 days of ICU hospitalization and SDD treatment. The carbapenemase *bla_KPC_* was detected in a single patient (patient #180), but only in the first sample (collected after 5 days in the ICU) and not in the second sample, which was collected after 16 days of ICU hospitalization. No other ESBL-or carbapenemase-producing strains were isolated from the patients during ICU hospitalization. Other Enterobacterial β-lactamases were found to be widespread in our resistome analysis. The *bla_AMPC_* β-lactamase was present in 37% of samples, with nine of ten patients and eight of ten healthy subjects having detectable levels of *bla_AMPC_* at one or more sampling points. The *bla_TEM_* β-lactamase was present in 26% of samples, corresponding with five of ten patients and four of ten healthy subjects in which this gene was detectable at one or more sampling points. None of the samples were positive for the carbapenemases *bla_NDM_* and *bla_OXA_*, or the transferable colistin resistance gene *mcr-1* (data not shown). Among resistance genes that are associated with Gram-positive pathogens, the staphylococcal methicillin resistance gene *mecA* was detected in 13 samples from eight of ten patients, but not in samples of healthy subjects. The vancomycin resistance gene *vanB* was present in 5 samples from three of ten patients and six samples from four of ten healthy subjects.

A comparison of the abundance of individual ARGs in samples that were collected during ICU stay, versus samples from healthy subjects, revealed that four ARGs (*aac(6′)-Ii*, *ermC*, *qacA*, *tetQ*) were significantly more abundant in ICU patients, while two ARGs (*catA* and *tetW*) were significantly more abundant in healthy individuals (Fig. 3).

**Figure 3.**
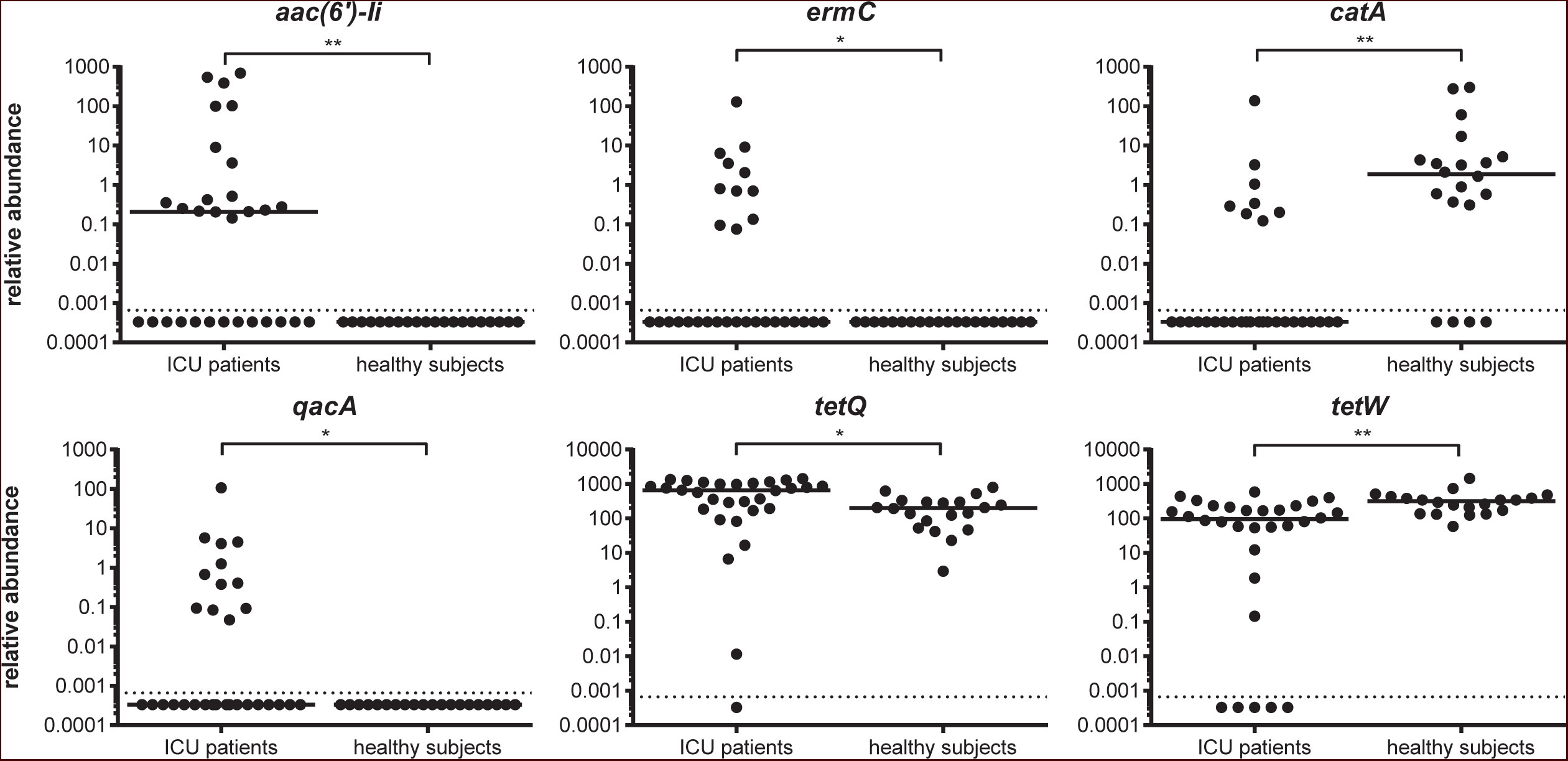
Antimicrobial resistance genes present at significantly higher or lower levels in the microbiota of ICU patients, compared to healthy subjects. ARGs that are present at significantly higher (*aac(6’)-Ii*, *ermC*, *qacA*, and *tetQ*) or lower (*catA* and *tetW*) abundance in ICU patients, compared to healthy subjects, are shown. Testing for statistically significant differences was performed by the Mann-Whitney U test, with Benjamini-Hochberg correction for multiple testing (* = *q* < 0.05; ** = *q* < 0.01). The horizontal line denotes the median value. The detection limit of the qPCR assay is indicated with the dashed line.

## Discussion

Current guidelines in the Netherlands recommend topical antibiotic decontamination in ICU patients with an expected ICU stay of two days or longer. Yet, the original claim that these interventions do not affect harmless anaerobic intestinal bacteria [7], has recently been questioned [20,34]. While culture-based studies did not demonstrate selection for antibiotic-resistant opportunistic pathogens during SDD-treatment [9,12,35,36], concerns remain that selection for antibiotic resistance genes occurs in the gut microbiota of patients.

The current study describes the dynamics of the gut microbiota of ICU-patients receiving SDD during ICU-stay towards dysbiosis, i.e. the perturbation of the complex commensal communities of the gut microbiota of healthy humans [37]. The gut microbiota of ICU patients was characterized by a low diversity, the increased abundance of enterococci and lower abundance of anaerobic Gram-positive, butyrate-producing bacteria of the *Clostridium* clusters IV and XIVa. These findings expand on previous findings of selection for Gram-positive cocci [12,20,38,39] and depletion of *F. prausnitzii* during SDD [34]. In addition, we were able to demonstrate that the abundance of *E. coli* was significantly lower in ICU-patients than in healthy individuals. The suppression of *E. coli* in the SDD-treated ICU patients starkly contrasts with other studies in critically ill patients not receiving SDD, in which high-level *E. coli* gut colonization is a common event [2,3,5]. This observation further supports previous studies which found that SDD is successful in suppressing outgrowth of *E. coli* in the gut microbiota of ICU patients [8,9], corresponding to the original aim of SDD [7].

Notably, levels of *E. coli* increased again after ICU-discharge in two of patients, reaching levels in the gut similar to, or even surpassing, those in healthy individuals. These findings suggest that a rapid regrowth or recolonization of the intestinal tract by *E. coli*, and possibly other aerobic Gram-negative bacteria, occurs upon cessation of prophylactic antibiotic therapy. It remains to be determined whether rapid post-ICU recolonization by *E. coli* increases the risk for infections with this bacterium. Yet, in the only prospective evaluation on the post-ICU effects of SDD, the implementation of SDD was not associated with higher infection rates after ICU discharge [40].

The qPCR-based analysis of the resistome confirms previous metagenomic studies, in showing that tetracycline and aminoglycoside resistance genes and bacteroidal β-lactamases are widespread in the human intestinal microbiota [14,15,18,20]. Four resistance genes (*aac(6′)-Ii*, *ermC*, *qacA*, *tetQ*) were significantly more abundant among ICU patients than in healthy subjects. The *aac(6’)-Ii* gene is a specific chromosomal marker for the nosocomial pathogen *Enterococcus faecium* and provides low-level resistance to aminoglycosides [41]. Its high abundance in ICU patients is in line with the increased levels of enterococci in the microbiota of the patients. The increased abundance of the macrolide resistance gene *ermC* may have been selected for by the use of low doses of the macrolide erythromycin, which was used as an agent to accelerate gastric emptying during ICU stay in six patients. The increased abundance of *tetQ* in the gut microbiota of ICU patients may reflect the higher abundance of Bacteroidetes in ICU patients versus healthy subjects, as *tetQ* is widely distributed on conjugative transposons in this phylum [42]. Finally, the *qacA* gene confers resistance to a number of disinfectants, including the biguanidine compound chlorhexidine and the quaternary ammonium compound benzalkonium chloride [43,44]. Disinfectants are widely used in ICUs as cleaning and infection control agents [45] and their use could select for *qacA* in the gut microbiota of patients.

Two resistance genes (*catA* and *tetW*) were more abundant in healthy individuals than in ICU patients. There is currently little information on the distribution of the *catA* gene among bacteria associated with the human gut microbiota, but the gene was frequently found in human faeces in a recent study set in low-income human habitats [46]. The tetracycline resistance gene *tetW* is present in Gram-positive anaerobic gut commensals [47]. The gut bacteria that *catA* and *tetW* are associated with, most likely members of *Clostridium* clusters IV and XIVa, are likely depleted during ICU-stay and SDD-treatment.

Although SDD improves survival of ICU-patients, its use remains controversial due to the perceived risk for selection of antibiotic resistance among bacteria that populate the patient gut. In this study, we were not able to include an ICU control group that was not treated with SDD, as this would be a breach of clinical guidelines for ICU-patients in our country. It is notable, however, that we did not find selection for high-risk antibiotic resistance genes (like ESBLs, carbapenemases or vancomycin resistance genes) in SDD-treated patients. The increased abundance of the resistance genes *aac(6’)-Ii*, *ermC, qacA* and *tetQ* in SDD-treated ICU patients in our study is – in our opinion - of limited concern. The first three resistance genes contribute to resistance in enterococci, either to relatively low concentrations of antibiotics (*aac(6’)-Ii*) or to classes of antimicrobials that are of limited relevance for the treatment of enterococcal infections (*ermC* and *qacA*). The *tetQ* gene provides resistance to tetracyclines in Bacteroidetes, but this class of antibiotics is scarcely used for the treatment of anaerobic infections [48]. Our data further support the ecological safety of SDD, at least in settings with low levels of circulating antibiotic resistance, as it does not lead to selection for clinically relevant antibiotic resistance phenotypes [36].

In addition, our study illustrates the expediency of using culture-independent methods to monitor the presence and abundance of antibiotic resistance genes. The qPCR platform used here enables the rapid detection and quantification of antibiotic resistance genes in the gut microbiota of patients. The detection of high-risk antibiotic resistance genes (encoding e.g. ESBLs, carbapenemases or vancomycin resistance proteins) in the resistome of patients may lead to the implementation of targeted antibiotic therapy or infection control measures to minimize the risk for selection and spread of these resistance genes.

## Declarations

### Ethics approval and consent to participate

The protocol for the ICU patient arm of this study was reviewed and approved by the institutional review board of the University Medical Center Utrecht (Utrecht, The Netherlands) under number 10/0225. Informed consent for faecal sampling during hospitalization was waived. The protocol for the feces collection of healthy subjects, including informed consent, was reviewed and approved by the Ethics Committee of Gelderse Vallei Hospital (Ede, The Netherlands).

### Consent for publication

Not applicable

### Availability of data and material

The datasets used and/or analysed during the current study available from the corresponding author on request.

### Competing interests

W.v. S. is a consultant for Vedanta Biosciences.

### Funding

This work was supported by The Netherlands Organisation for Health Research and Development ZonMw (Priority Medicine Antimicrobial Resistance; grant 205100015) and by the European Union Seventh Framework Programme (FP7-HEALTH-2011-single-stage) ‘Evolution and Transfer of Antibiotic Resistance’ (EvoTAR), under grant agreement number 282004. In addition, W.v.S is is supported by a NWO-VIDI grant (917.13.357)

## Authors’ contributions

RJLW, MJMB, MWJvP, HS and WvS designed the study. EB, MSvM, EANO, MWJvP, HS and WvS supervised collection of fecal samples. EB, TdJBG, EAMM and JCB performed experiments to map the gut resistome and phylogenetic composition of the microbiome. SF, WAAdSP, LL and JRB contributed bioinformatic analyses. The manuscript was written by EB, TdJBG, SF, RJLW, MJMB, MWJvP, HS and WvS and approved by all authors.

## Acknowledgements

We thank ServiceXS B.V. (Leiden, The Netherlands) for their assistance in the Fluidigm real-time PCR assays. We are grateful to Erwin Zoetendal and Willem M. de Vos, providing material and data from the Cohort study of intestinal microbiota among Irritable Bowel Syndrome patients and healthy individuals’ (CO-MIC) funded by the unrestricted Spinoza Award to Willem M. de Vos from the Netherlands Organization for Scientific Research.

